# An extension of Shannon’s entropy to explain taxa diversity and human diseases

**DOI:** 10.1101/2020.08.03.233767

**Authors:** Farzin Kamari, Sina Dadmand

**Author notes:** **Corresponding author:** Farzin Kamari **Email:**.

## Abstract

In this study, with the use of the information theory, we have proposed and proved a mathematical theorem by which we argue the reason for the existence of human diseases. To introduce our theoretical frame of reference, first, we put forward a modification of Shannon’s entropy, computed for all available proteomes, as a tool to compare systems complexity and distinguish between the several levels of biological organizations. We establish a new approach to differentiate between several taxa and corroborate our findings through the latest tree of life. Furthermore, we found that human proteins with higher mutual information, derived from our theorem, are more prone to be involved in human diseases. We further discuss the dynamics of protein network stability and offer probable scenarios for the existence of human diseases and their varying occurrence rates. Moreover, we account for the reasoning behind our mathematical theorem and its biological inferences.

## Introduction

The term ‘entropy’ was originally introduced by Rudolf Clausius in thermodynamics more than one and a half centuries ago (Clausius, 1864). Entropy is predominantly known as a measure of the disorder and uncertainty in dynamic systems (Bailey, 2009; Ghahramani, 2006). In information theory, entropy, also known as Shannon’s entropy, is defined as the average minimum rate at which information is produced or predicted in an uncertain stochastic setting (Shannon, 1948). In recent decades, information theory has been vastly applied in many fields of science (Andrews et al., 2015). Biology is of no exception, but compared to other areas, the applications of information theory in biological sciences have been indeed limited (Battail, 2013). More importantly, medical sciences lack any use of information theory in daily practice. The applications of information theory in molecular biology have been mostly focused on genome sequence analysis (Vinga, 2013). To date, no study has investigated the evolutionary nature of human diseases using information theory.

The backbone of evolution is random genetic mutations being selected according to the natural environment. So what has been encountered in nature after some 3.5 billion years of life history is a ‘selected randomness’. This is the reason why we believe information theory can be a perfect language to understand life – i.e., this selected randomness. In the literature, single nucleotide polymorphisms (SNPs) accounting for the main portion of this randomness have been associated with inherited disease susceptibility (Bodmer & Bonilla, 2008; Z. Wang & Moult, 2001). However, such approaches have only focused on the genome investigation and, in most part, neglected the human proteome and the protein-protein interactions (PPIs). PPIs are the leading cause of cellular metabolic processes. They are induced-fit physical contacts between macromolecules of proteins allowing the cellular function (Changeux & Edelstein, 2011; Keskin et al., 2008; Koshland Jr, 1995). In order to employ information theory in medical sciences, it would be necessary to investigate diseases in detail considering their molecular networks and PPIs. We believe evolutionary evidence interpreted by stochastic information analysis can provide substantial help in understanding diseases of living organisms.

In this study, to understand the nature of human diseases, we have focused on human interactome and available proteomes of living organisms. To avoid confusion, by ‘human diseases’ we only refer to non-communicable diseases in human with at least one reported genetic basis. Also, as the term ‘proteome’ has sometimes referred to all proteins of a cell or a tissue in the literature, it is to be noted that in this article, the term ‘proteome’ will refer to the complete set of proteins that *can be* expressed by an *organism*. Because proteomes are functional representatives of the ‘expressed genome’ of organisms, we have used them as the means of our investigation. We have used Shannon’s entropy as a retrograde approach to trace ~180 million proteins with more than 61 billion amino acids through the tree of life and investigated the trends of complexity among organisms. We have shown that this methodology agrees with the classification of phyla and may be used as a new tool in taxonomy. Also, using our new mathematical theorem presented in the Methods section, we have focused on *Homo sapiens’* PPI network and discussed potential clinical applications in the practice of medicine. We argue why there are only the diseases we know, and not others, and discuss why some diseases are more prevalent. We also elaborate on the reasonable links between our mathematical theory, Shannon’s entropy, the evolution of taxa, and human diseases.

## Results

### CAIR comparisons among taxonomic groups

Calculated Average Information per Residue (CAIR) was calculated (see Methods) for all proteomes available at the UniProt database until April 2020. Nearly 180 million proteins with more than 61 billion amino acids were analysed to classify ~29,000 organisms in 92 phyla. Table 1 shows CAIRs of the most popular proteomes and model organisms (for all ~29k proteomes, see Supplementary Table 1). The minimum CAIR of an organism is that of *Zinderia insecticola* (0.8247) and the maximum is of *Ciona savignyi* (0.9449). The mean ± standard deviation considering all organisms is 0.9210 ± 0.0129 with a median (interquartile range) of 0.9250 (0.0160). Having performed a literature review of articles published no later than April 2020, we have drawn the most updated tree of life for UniProt taxonomic lineage data (Figure 1A). For each bifurcation point on the tree, we tested if two sides of the bifurcation have developed divergent CAIRs. Figure 1B illustrates how CAIR divergence is present through the different lineages of taxonomy. On all bifurcation points of Figure 1A, a number is written whose respective statistical test results are demonstrated in Figure 1B with two half violin plots for upper and lower sides of the bifurcation and their box-and-whisker plots. Since the groups were negatively skewed, unbalanced, and heteroscedastic, their difference was investigated via the two-sided Brunner-Munzel statistical test (Neuhäuser & Ruxton, 2009). It is noteworthy that groups with ten or fewer organisms were excluded from comparisons, as the Brunner-Munzel test is statistically imprecise even with a permutation. Among 56 performed tests, 48 tests demonstrated a significant difference at the point of bifurcation. Interestingly, the bifurcation points of eight insignificant tests are mostly known to be a matter of controversy in the scientific literature (Evans et al., 2019; Spang et al., 2017). Along with the *p*-value significance, estimated effect sizes (ES) and 95% confidence intervals (CI) are also reported. For the exact *p*-values of each test, please see Table 2.

**Figure 1.**
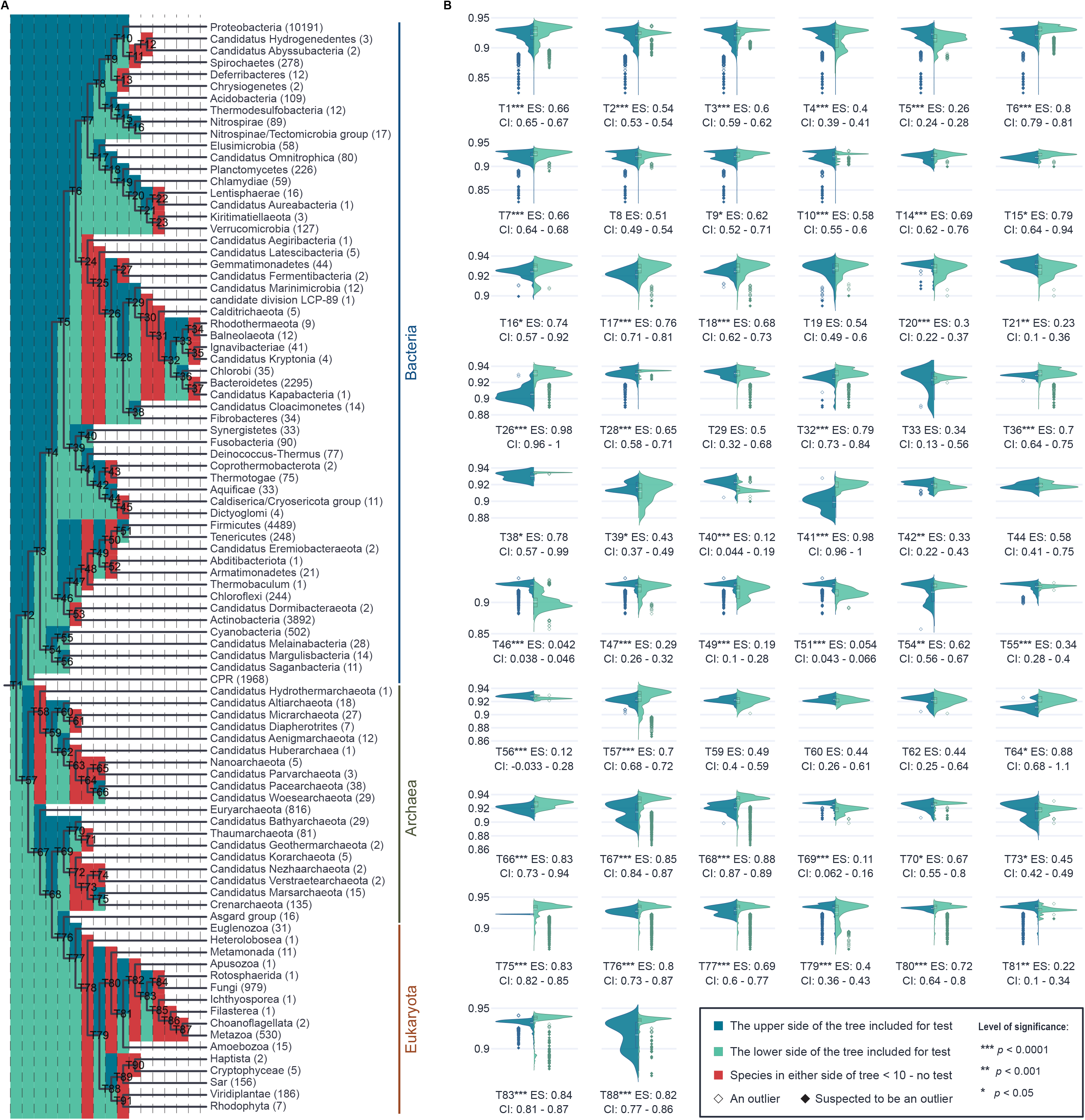
CAIR comparisons through the tree of life. **(A)** The most updated tree of life comprising all second hierarchies stemming from cellular organisms. For bacteria superkingdom, the second hierarchy includes all 56 bacterial phyla; Candidate Phyla Radiation (CPR) is included as one separate phylum. For Archaea and Eukaryota superkingdoms, the second hierarchy encompasses 20 archaeal phyla and 16 eukaryote supergroups and divisions. On each bifurcation point of the tree, the arbitrary number corresponds to the test number and the associated plots in panel B. The blue and light green colours indicate the superior and inferior arm of the bifurcation point, respectively, which correspond to the left and right sides of the violin- and box-and-whisker plots. The red colour indicates a bifurcation point with both arms from which at least one arm contains less than ten organisms and thus the Brunner-Munzel test would not be reliable. The numbers in parentheses designate the total number of available complete proteomes in each group. **(B)** Violin plots of each bifurcation point in the tree of life, except for those in red. The vertical axes refer to the CAIR in all plots. The box-and-whiskers are overlaid within each violin plot, and the white dashed line in each box indicates the CAIR mean in the corresponding group of organisms. None of the outliers were excluded from the analysis. Asterisks after each test number indicate the significance level of tests. ES; estimated effect size. CI; 95% confidence interval.

**Table 1.**
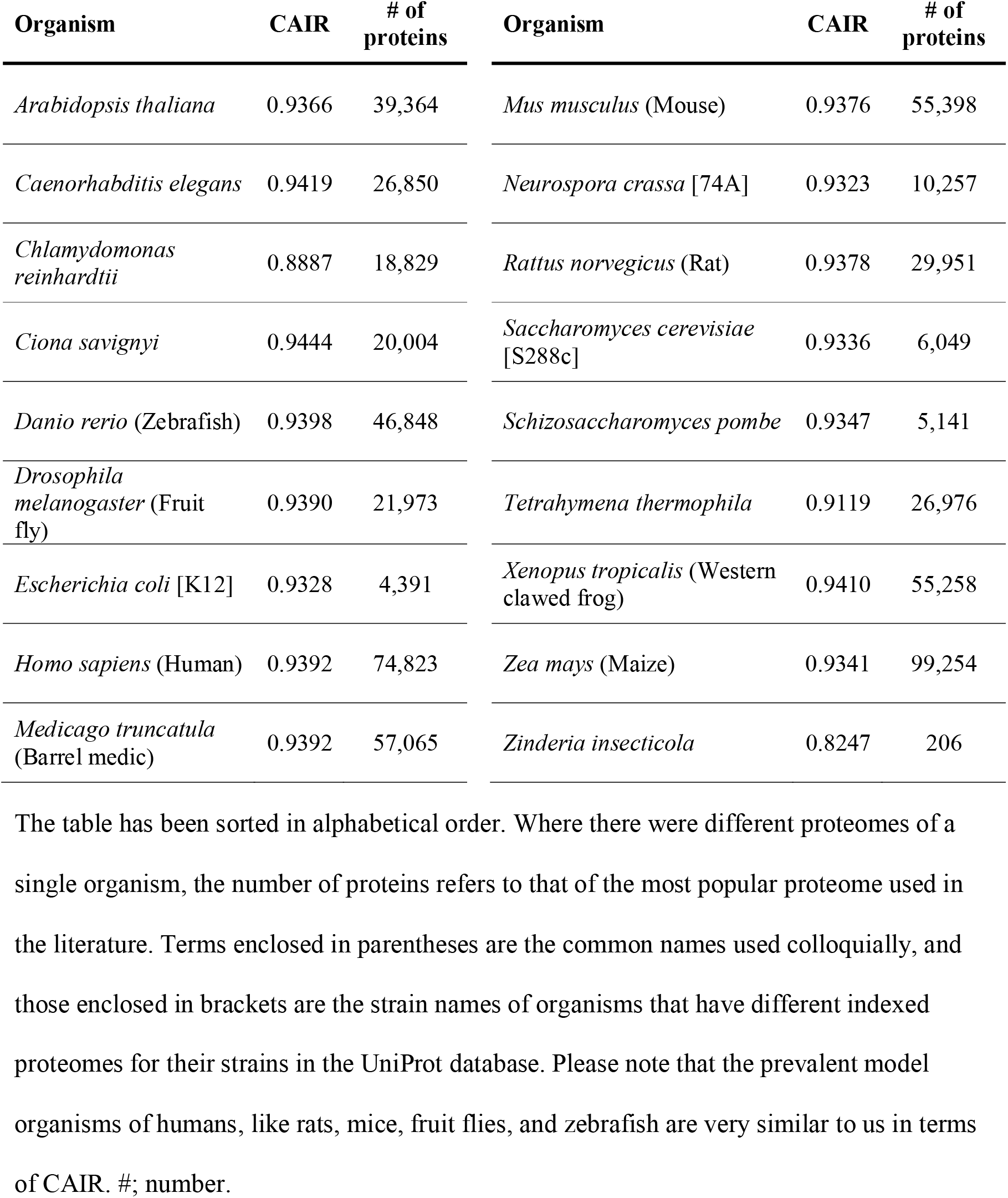
Proteome CAIRs of organisms mainly used in biological models and studies.

**Table 2.** Exact *p*-values of the two-sided Brunner-Munzel tests.

| Test no. | $p$-value | Test no. | $p$-value | Test no. | $p$-value | Test no. | $p$-value |
| --- | --- | --- | --- | --- | --- | --- | --- |
| T1 | $5 \times 10^{-137}$ | T18 | $2.72 \times 10^{-10}$ | T42 | 0.000965 | T66 | $4.29 \times 10^{-08}$ |
| T2 | $3.42 \times 10^{-17}$ | T19 | 0.14649 | T44 | 0.35037 | T67 | 0 |
| T3 | $6.91 \times 10^{-31}$ | T20 | $6.98 \times 10^{-07}$ | T46 | 0 | T68 | 0 |
| T4 | $7.42 \times 10^{-131}$ | T21 | 0.000417 | T47 | $2.88 \times 10^{-28}$ | T69 | $5.64 \times 10^{-32}$ |
| T5 | $2.92 \times 10^{-76}$ | T26 | $2.74 \times 10^{-42}$ | T49 | $2.51 \times 10^{-07}$ | T70 | 0.008022 |
| T6 | 0 | T28 | $1.66 \times 10^{-05}$ | T51 | $7.23 \times 10^{-205}$ | T73 | 0.004432 |
| T7 | $6.41 \times 10^{-47}$ | T29 | 0.99445 | T54 | 0.000363 | T75 | $2.71 \times 10^{-263}$ |
| T8 | 0.32303 | T32 | $8.86 \times 10^{-17}$ | T55 | $5.04 \times 10^{-07}$ | T76 | $7.62 \times 10^{-08}$ |
| T9 | 0.019045 | T33 | 0.14933 | T56 | $5.13 \times 10^{-05}$ | T77 | $9.92 \times 10^{-05}$ |
| T10 | $6.98 \times 10^{-08}$ | T36 | $3.83 \times 10^{-09}$ | T57 | $8.48 \times 10^{-56}$ | T79 | $1.30 \times 10^{-08}$ |
| T14 | $7.70 \times 10^{-08}$ | T38 | 0.012529 | T59 | 0.87909 | T80 | $7.16 \times 10^{-05}$ |
| T15 | 0.001135 | T39 | 0.030999 | T60 | 0.46556 | T81 | 0.000134 |
| T16 | 0.010167 | T40 | $5.65 \times 10^{-16}$ | T62 | 0.55498 | T83 | $6.78 \times 10^{-89}$ |
| T17 | $2.45 \times 10^{-17}$ | T41 | $2.73 \times 10^{-51}$ | T64 | 0.002841 | T88 | $1.35 \times 10^{-34}$ |

### CAIR as a means of understanding the behaviour of natural selection

The extent of natural selection’s capacity to show bias in favour of selecting a spectrum of organisms is open to question. Since genetic mutations are known to be generally random, the CAIR density plot of such a random condition without a selection bias shall turn out to have a uniform distribution. In a simulation of various random protein systems, we obtained a similar distribution to the actual CAIR density plot taking into account a negative skewness of −0.90 (Figure 2). Figure 2A shows the density plot of life and Figure 2B depicts that of our simulation. Figure 2C, also, shows how these two distributions are similar given a tiny bandwidth. To test for their similarity of distributions, we also performed two-sample Kolmogorov-Smirnov tests whose mean *p*-value was 0.40 on 1000 iterations. It is noteworthy, not unexpected though, that the natural selection is biased toward the organisms with higher CAIRs. This might have stemmed from the random mutations over the course of ages as discussed in the next section. Also, natural selection favours more complex and more unpredictable protein systems, as they can accommodate superior functionalities. Besides, the density plot of CAIRs possesses one other interesting property. Since there are significant differences among taxonomic groups of the second hierarchy as shown in Figure 1B, it can further be expected that all organisms are noticeable on the density plot. Figure 3 shows how different taxonomic hierarchies are manifested in the density plot of organism CAIRs. On the ‘wave of life’, members of the succeeding taxonomic ranks are revealed by zooming in on the preceding taxonomic group. This property of CAIR density suggests an original methodology to help classifying organisms into various taxa.

**Figure 2.**
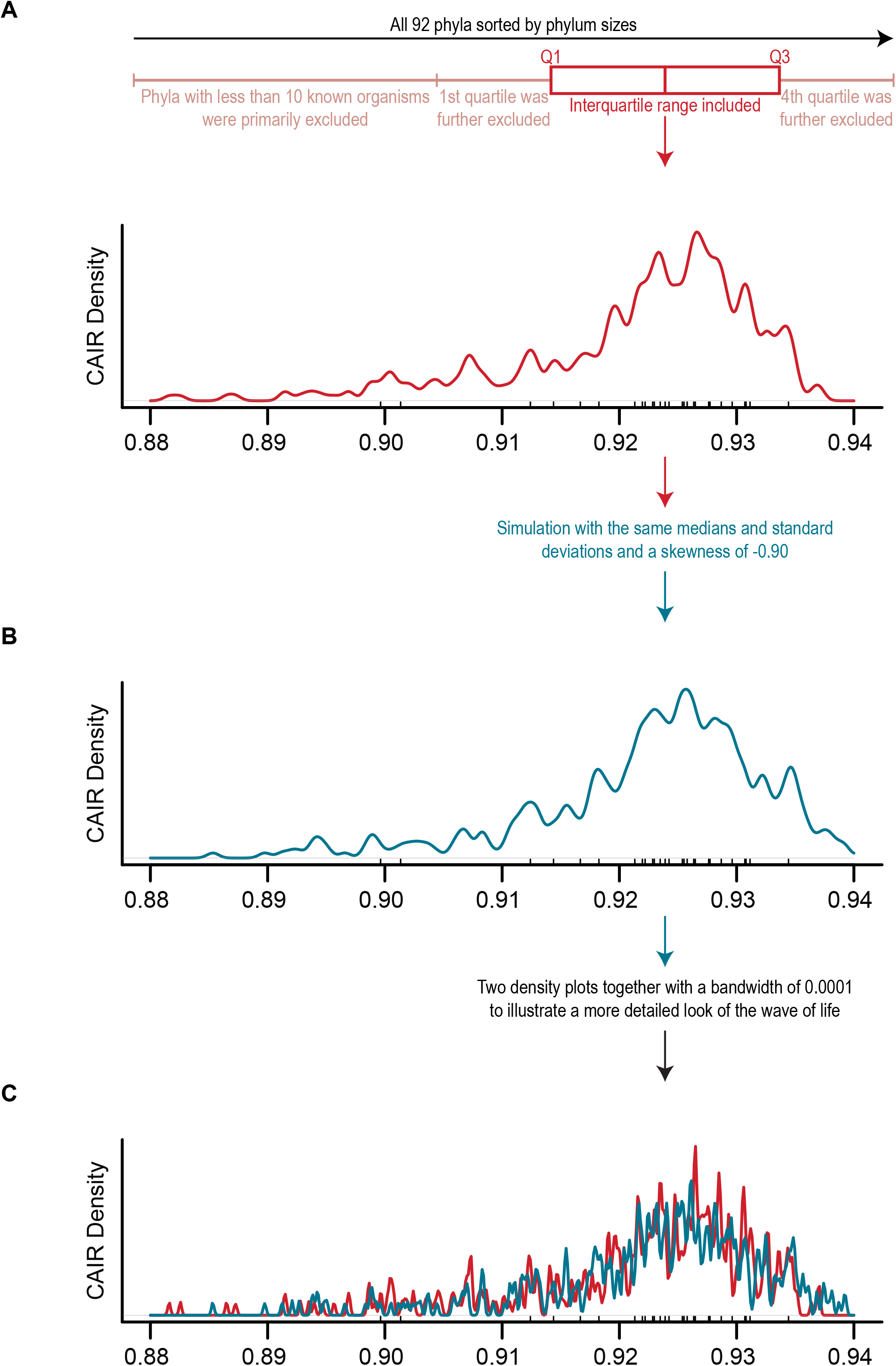
A simulation of natural selection shows the selection’s bias towards higher CAIRs. **(A)** CAIR density plot of the organisms considering the interquartile range (IQR) of phyla sizes through the tree of life after removing the tiny phyla with less than 10 organisms (Q_1_-Q_3_ 16.5 – 171). From the 27 phyla within the IQR, random sampling was performed with a size of Q_1_ ~ 17. IQR inclusion and the subsequent sampling was done to remove the size effects of populated phyla on the density plot. The mean of phyla skewness is −0.82 in Q_0_-Q_4_ and −0.74 in Q_1_-Q_3_. **(B)** CAIR simulation of the tree of life with 27 negatively skewed normal distributions. The means of these simulated random distributions equal to the respective medians of the 27 phyla (marginal rugs) explained in panel A. The function was iterated 1000 times to find the skewness in which the Kolmogorov-Smirnov (KS) test has the maximum *p*-value. The simulation revealed a skewness of −0.90. **(C)** Both panels of A and B are overlaid with a lesser bandwidth to show the details of the distributions. KS test reveals a *p*-value of 0.40 not rejecting the null hypothesis that distributions are identical. Negative skewness of the red wave (depicting phyla data) suggests that the natural selection is biased in favour of organisms with higher CAIRs. Q; quartile.

**Figure 3.**
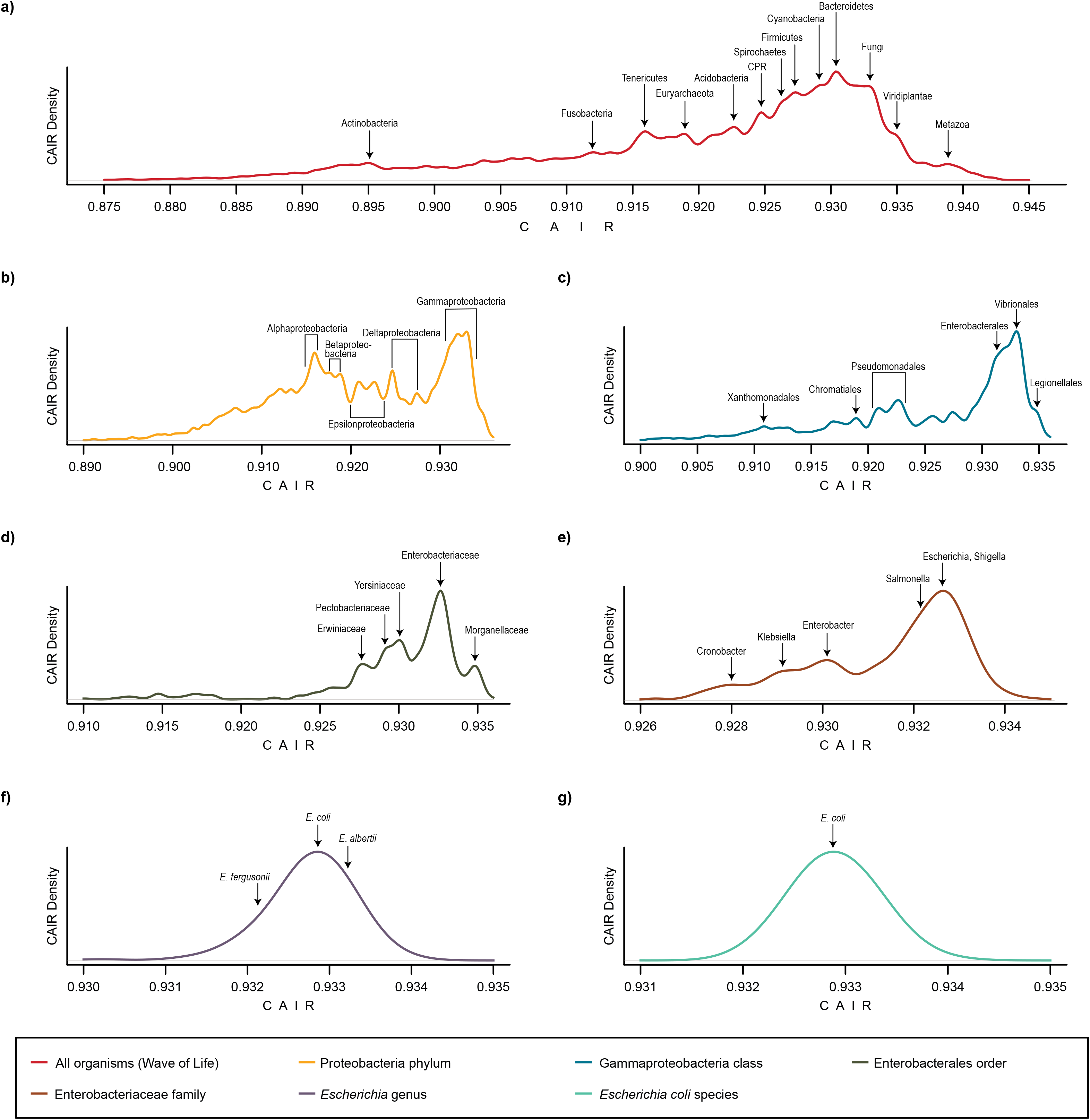
From the whole life to *E. coli* as an exemplary organism illustrating the CAIR density of proteomes in several ranks of taxonomy. **(A)** The ‘wave of life’ denotes the CAIR density of proteomes through the tree of life. As it is argued, the wave of life is proposed to be a summation of skewed distributions. As a result, the wave of life holds an interesting property; namely, zooming in on the wave of life by narrowing the range on the horizontal axis reveals the members in the next rank of taxonomy. In all panels, the peaks of the plots are named according to the most abundant phyla with the closest median to the peak. **(B)** CAIR density of the proteomes in the Proteobacteria phylum. **(C)** Zooming in further on panel B and narrowing the range of horizontal axis to the distribution of organisms under Gammaproteobacteria class, i.e. CAIRs of 0.900 – 0.936, reveals the taxonomic orders. **(D-G)** Proceeding further to zoom in on the peaks of the previous plot shows a perfect agreement with all taxa lineage. Lastly, several strains of *E. Coli* form a bell-shaped distribution, as shown in panel G.

### Human proteome analysis and the estimation of mutual information for a protein (EMIP)

According to the theorem presented in the Methods section and its biological inferences, EMIP has been calculated for each human protein entry using its PPI network. Each Swiss-Prot, i.e. reviewed, entry has been categorized into three groups using the Orphanet database of diseases. In case of a reported disease(s) related to an entry, all disease point-prevalence/incidences of the entry (from Orphanet database) are summed up to obtain the total occurrence of the disease, that is to say, the protein’s overall malfunction. Table 3 shows the narrative data of the human disease categories. As seen in the table, the groups are unbalanced in size and heteroscedastic which makes conventional statistical analyses unfavourable. Herein, our results demonstrate how well indicators of diseases can be correlated to the disease occurrence categories. In the Methods section, we have explained why such independent variables were candidates of correlation and further statistical analyses. Figure 4 shows the results of comparisons between disease occurrence categories in four disease indicators. In each comparison, we have also included gene age categories to test our hypotheses and biological inferences. The Dunnett-Tukey-Kramer pairwise multiple comparison test adjusted for unequal variances and unequal sample sizes (Dunnett, 1980) was performed to test overall comparisons. Results of comparisons show that disease indicators correlate significantly with disease occurrence categories. Among four indicators, EMIP is revealed to have the most significant differences between categories, while CAIR was incongruous. The inconsistency seen in CAIR confirms that the use of Shannon’s entropy alone is not a good enough indicator of disease occurrence categories.

**Figure 4.**
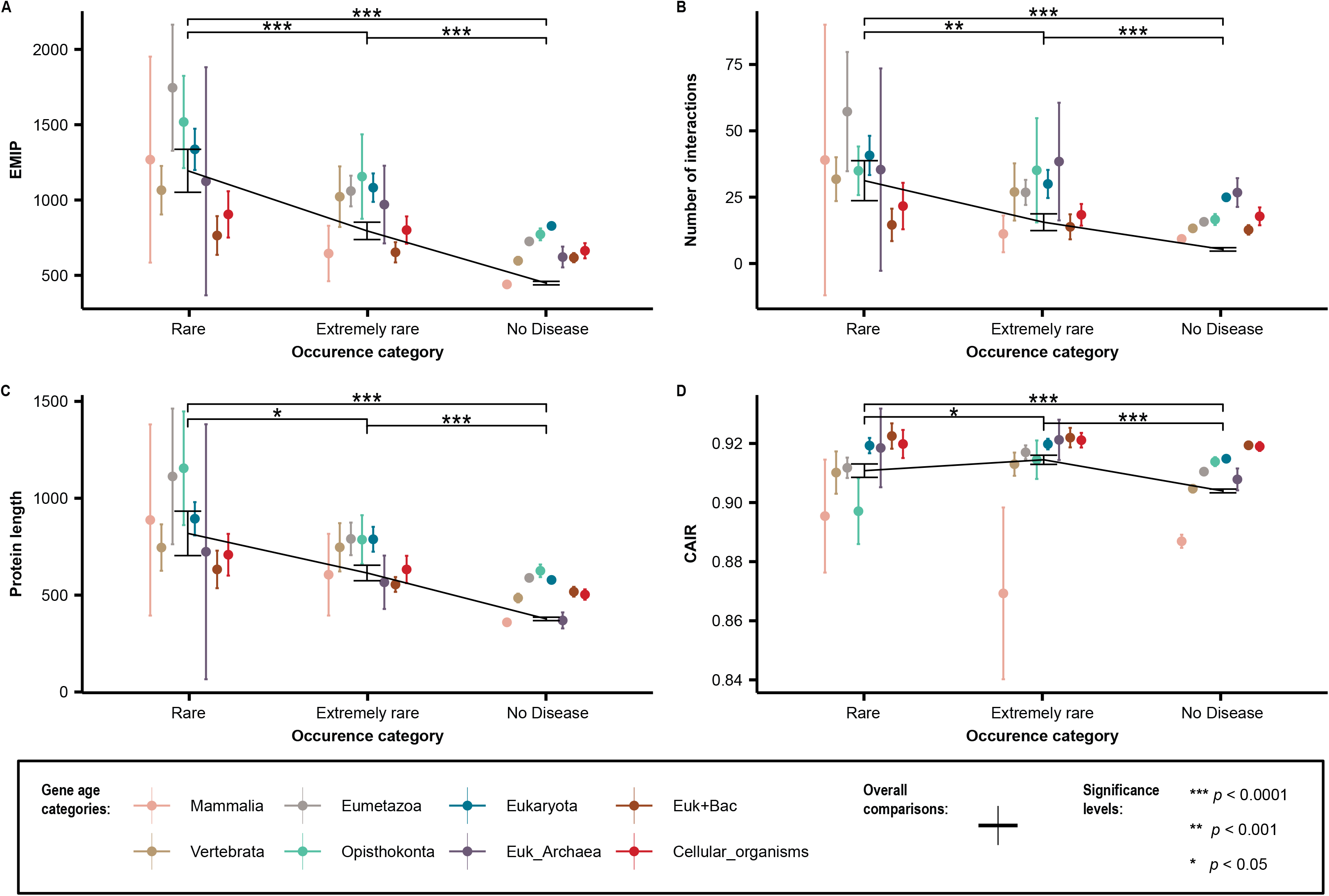
Comparisons of disease indicators in three occurrence groups across gene age categories. **(A)** EMIP increases significantly as the occurrence of diseases increases. Generally, EMIP has the highest level of significance among disease indicators. Not surprisingly, the trend of EMIP is also increasing as the genes age. The decline of EMIP in primordial gene age eras are due to the eukaryotic branching and the nucleic genetic material. **(B)** The number of interactions is the second-best indicator of the disease occurrence category. Similarly, as expected, the trend of interactions is increasing as genes grow older. Evolution brings new interactions and adds new nodes to the network. Similarly, there is a decline in the number of interactions in the last two gene age eras. **(C)** The bigger the size of a protein, the more likely it is to be involved in a disease. Also, it is noticeable that the gene ages correlates positively with the protein size in both eukaryotic and prokaryotic settings. However, the disparity between these two settings is easily discernible. **(D)** Unlike what is expected, extremely rare diseases account for the proteins with the highest CAIRs. This observation has been further elucidated in the Discussion section. Nonetheless, the trend of gene ages agrees with the expectation as the complexity of proteins increases with age. Error bars illustrate mean ± 95% confidence intervals, and the significance test is the Dunnett-Tukey-Kramer pairwise multiple comparison test adjusted for unequal variances and unequal sample sizes.

**Table 3.**
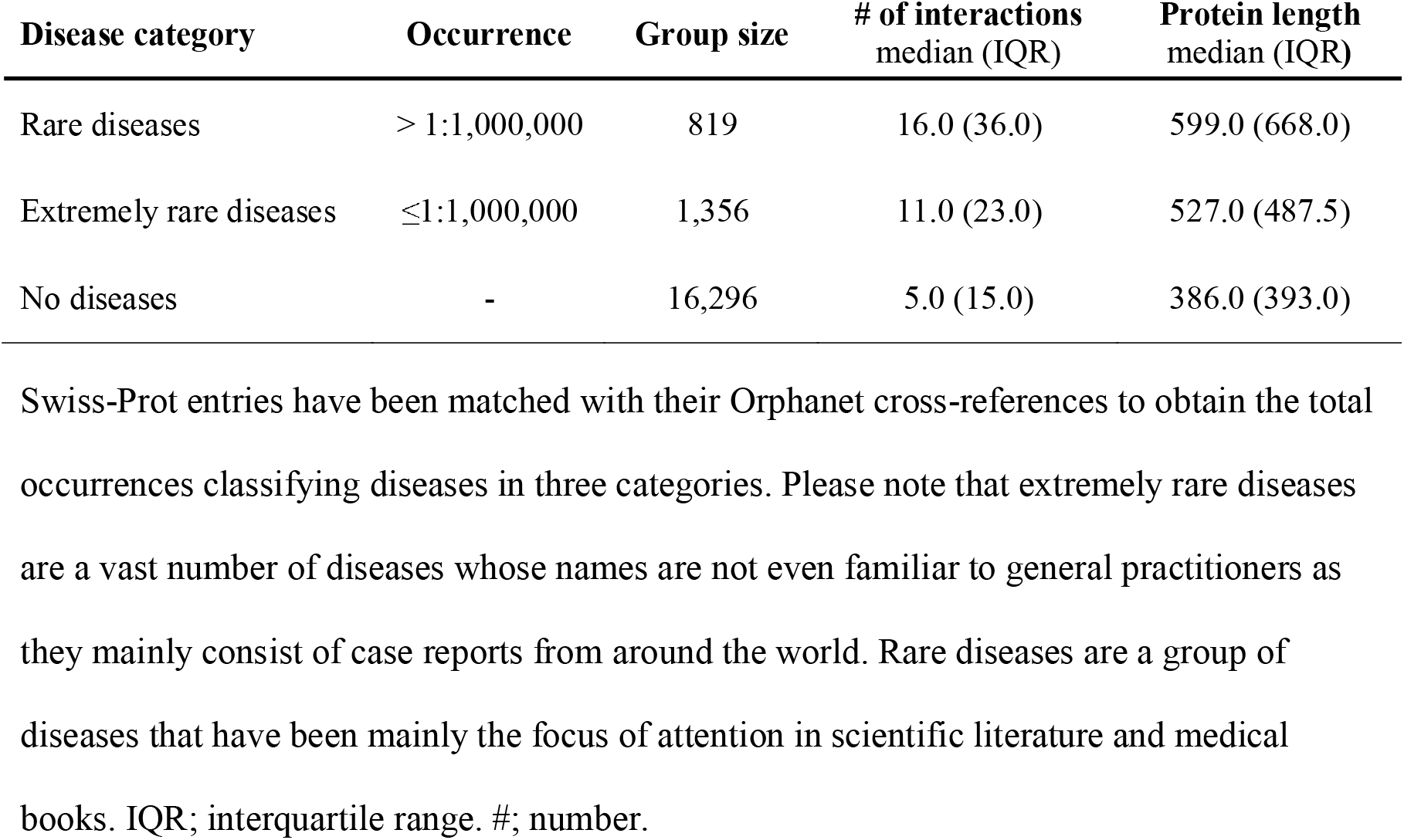
Narrative data of protein groups in three human disease categories.

| **Disease category** | **Occurrence** | **Group size** | **# of interactions**<br>**median (IQR)** | **Protein length**<br>**median (IQR)** |
| --- | --- | --- | --- | --- |
| Rare diseases | > 1:1,000,000 | 819 | 16.0 (36.0) | 599.0 (668.0) |
| Extremely rare diseases | ≤1:1,000,000 | 1,356 | 11.0 (23.0) | 527.0 (487.5) |
| No diseases | - | 16,296 | 5.0 (15.0) | 386.0 (393.0) |

Swiss-Prot entries have been matched with their Orphanet cross-references to obtain the total occurrences classifying diseases in three categories. Please note that extremely rare diseases are a vast number of diseases whose names are not even familiar to general practitioners as they mainly consist of case reports from around the world. Rare diseases are a group of diseases that have been mainly the focus of attention in scientific literature and medical books. IQR; interquartile range. #; number.

Additionally, since gene ages are presented as ranked data in the literature (Liebeskind et al., 2016), ranked analysis with an equal number of ranks has been performed to make all five indicators comparable with one another. Figure 5 shows the Likert plots and rank comparisons. As illustrated, natural Logarithm of EMIP (LEMIP) is by far better than other indicators in correlating disease occurrence categories which might be stemmed from its bell-shaped histogram. Unlike other indicators, the distribution of LEMIP allows it to be ranked with mean and standard deviations which have been reported in the Table 4. Please refer to Table 4 for detailed information about the ranks in disease indicators.

**Figure 5.**
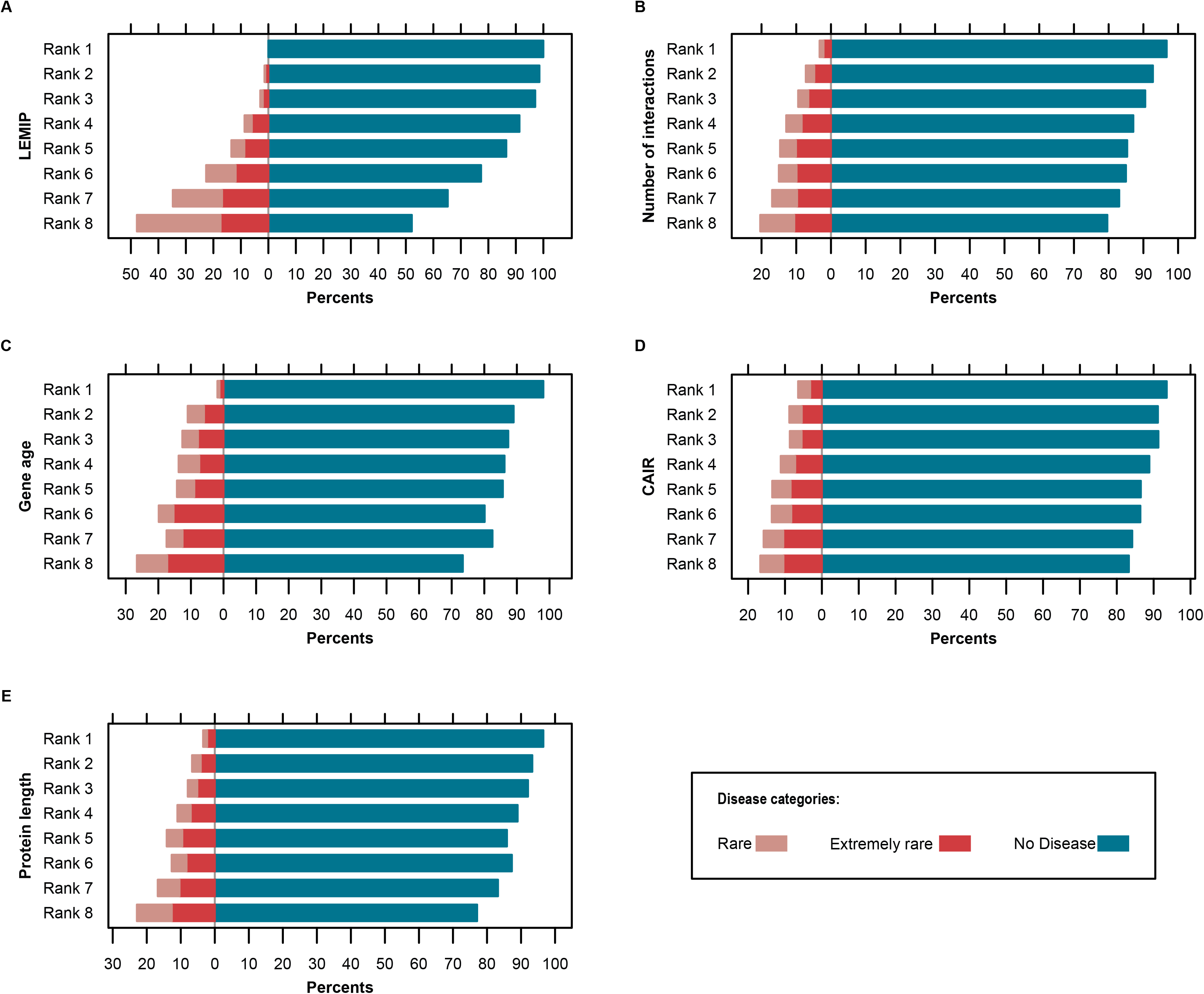
Likert plots of disease indicators (all classified into 8 ranks) with regard to occurrence categories. **(A)** Ranked data of EMIP show a robust relationship with the disease categories. Note that the occurrence of diseases increases as EMIP increases. Because the log-transformed of EMIP (LEMIP) forms a bell-shaped curve, the ranking has been done by the mean and three standard deviations of LEMIP. **(B)** Ranked data of interactions increase with the occurrences, maintaining the order of ranks. However, the number of interactions is not as well correlated to occurrences as EMIP. It is noteworthy in B, D, and E panels, since indicator distributions are inconsistent with a Gaussian distribution, rankings have been accomplished by median and equal percentile intervals. **(C)** The link between gene age ranks and disease categories is satisfactory considering the first and last ranks; however, the overall order of ranks does not match with the order of occurrence categories. Gene age ranks are the same as the eight age groups presented previously in the literature (Liebeskind et al., 2016). **(D)** Among Likert plots, CAIR ranks have shown the least correlation with the disease categories which is in line with Figure 4D. **(E)** The ranks of protein length are associated with disease occurrence categories maintaining the order of ranks, except for the sixth rank.

**Table 4.** Eight ranks of disease indicators and their quantitative intervals.

| Disease indicator | Rank 1 | Rank 2 | Rank 3 | Rank 4 | Rank 5 | Rank 6 | Rank 7 | Rank 8 |
| --- | --- | --- | --- | --- | --- | --- | --- | --- |
| LEMIP<sup>a</sup> | [-9, 36.45] | (36.44, 44.94] | (44.94, 53.44] | (68.38, 61.94] | (61.94, 70.44] | (70.44, 78.93] | (78.93, 87.43] | (87.43, 104.00] |
| Number of interactions<sup>b</sup> | [0] | [1] | (1, 3] | (3, 6] | (6, 11] | (11, 19] | (19, 39] | (39, 2136] |
| Protein length<sup>b</sup> | [2, 161] | (161, 250] | (250, 328] | (328, 415] | (415, 518] | (518, 671] | (671, 968] | (968, 34350] |
| CAIR<sup>b</sup> | [0.0217, 0.8803] | (0.8803, 0.8982] | (0.8982, 0.9088] | (0.9088, 0.9164] | (0.9164, 0.9230] | (0.9230, 0.9290] | (0.9290, 0.9351] | (0.9351, 0.9567] |
| Gene age<sup>c</sup> | Mammalia | Vertebrata | Eumetazoa | Opisthokonta | Eukaryota | Euk_Archaea | Euk+Bac | Cellular_organisms |

a. Because of the bell-shaped curve distribution, mean and standard deviations have been used to classify ranks. b. Because of the non-normal distribution, median and percentiles have been used to classify ranks. c. Ranks are based on scientific literature. LEMIP; Log<sub>e</sub> of Estimation of Mutual Information of Proteins. CAIR; Calculated Average Information per Residue.

As results suggest, EMIP seems to be a superior indicator of human diseases. Calculated values of disease indicators for all reported diseases (Supplementary Table 2) and all human proteins (Supplementary Table 3) are available in the Supplementary Material. In a nutshell, we have presented 16 human proteins with the highest EMIPs in Table 5. It is noteworthy that high-EMIP proteins are more susceptible to have diseased networks and are clinically crucial for human health. Figure 6 shows the network topology of the same proteins (for its R code, see the Data Availability section).

**Figure 6.**
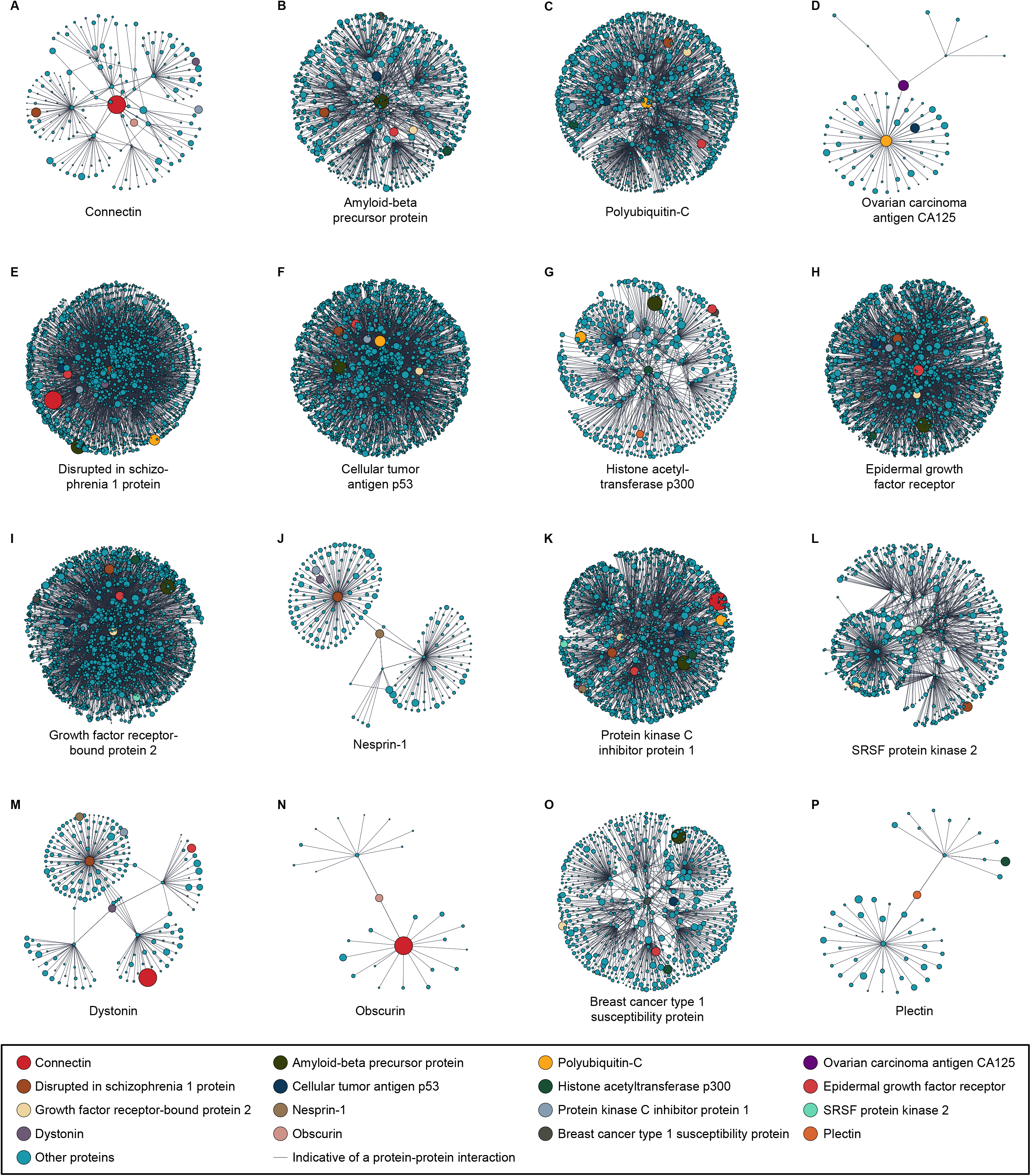
Protein networks of the proteins with the highest EMIPs. **(A-P)** The illustrated networks represent the proteins in Table 5 as the 16 proteins with the highest EMIPs in the human proteome. The size of each circle represents the EMIP of the protein. According to the UniProt database, networks illustrated in C, D, I, K, and L panels are non-disease networks; yet, the rest are involved in at least one disease. The main proteins are demonstrated in colours other than sky blue, and all other interactors are coloured in sky blue. The figure has been drawn with the interactions data available at the UniProt website up to the second-degree interactions. It is noteworthy that the illustrated proteins with the highest EMIP values are markedly present in various disease networks.

**Table 5.**
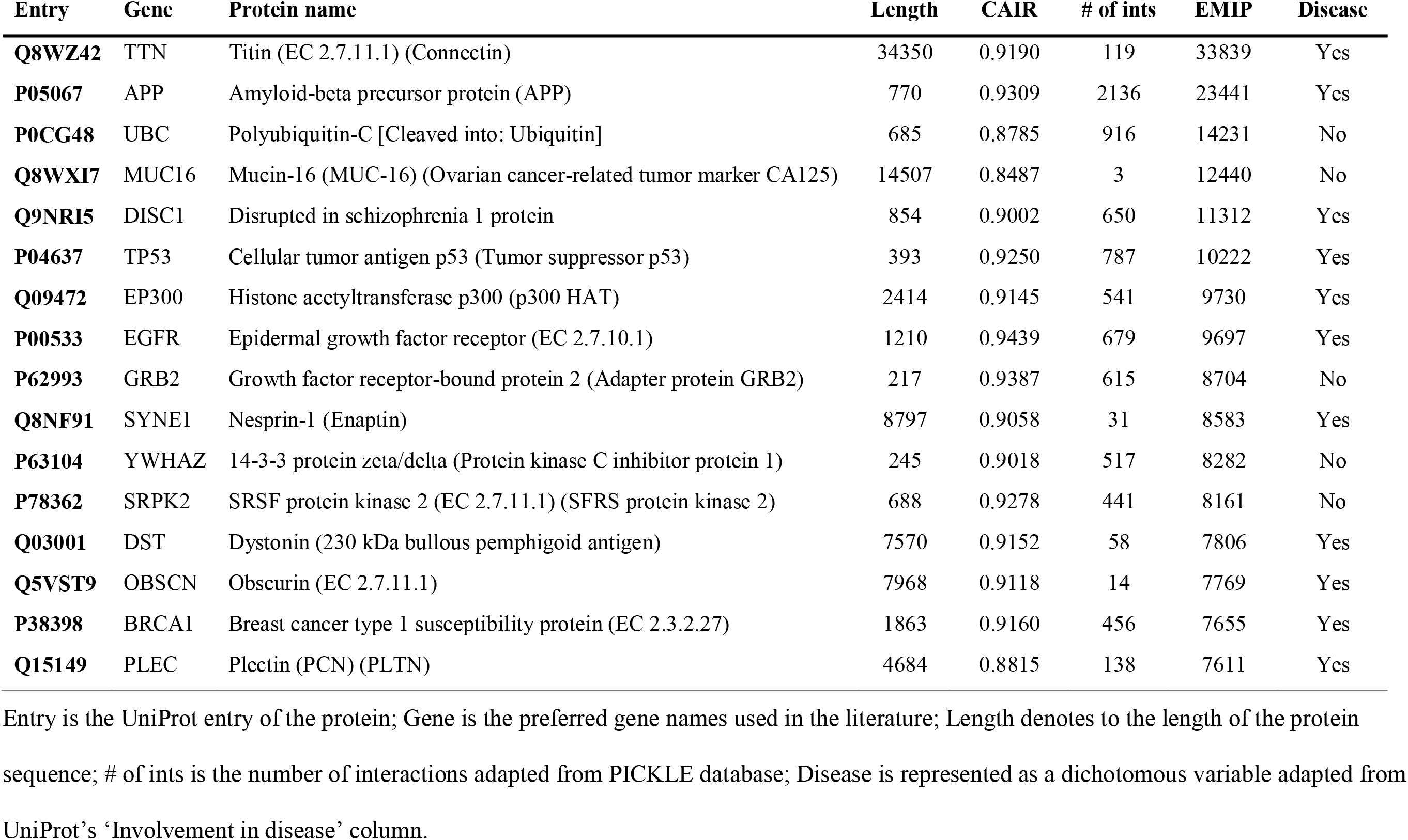
Proteins with the highest EMIP values.

| Entry | Gene | Protein name | Length | CAIR | # of ints | EMIP | Disease |
| --- | --- | --- | --- | --- | --- | --- | --- |
| **Q8WZ42** | TTN | Titin (EC 2.7.11.1) (Connectin) | 34350 | 0.9190 | 119 | 33839 | Yes |
| **P05067** | APP | Amyloid-beta precursor protein (APP) | 770 | 0.9309 | 2136 | 23441 | Yes |
| **P0CG48** | UBC | Polyubiquitin-C [Cleaved into: Ubiquitin] | 685 | 0.8785 | 916 | 14231 | No |
| **Q8WXI7** | MUC16 | Mucin-16 (MUC-16) (Ovarian cancer-related tumor marker CA125) | 14507 | 0.8487 | 3 | 12440 | No |
| **Q9NRI5** | DISC1 | Disrupted in schizophrenia 1 protein | 854 | 0.9002 | 650 | 11312 | Yes |
| **P04637** | TP53 | Cellular tumor antigen p53 (Tumor suppressor p53) | 393 | 0.9250 | 787 | 10222 | Yes |
| **Q09472** | EP300 | Histone acetyltransferase p300 (p300 HAT) | 2414 | 0.9145 | 541 | 9730 | Yes |
| **P00533** | EGFR | Epidermal growth factor receptor (EC 2.7.10.1) | 1210 | 0.9439 | 679 | 9697 | Yes |
| **P62993** | GRB2 | Growth factor receptor-bound protein 2 (Adapter protein GRB2) | 217 | 0.9387 | 615 | 8704 | No |
| **Q8NF91** | SYNE1 | Nesprin-1 (Enaptin) | 8797 | 0.9058 | 31 | 8583 | Yes |
| **P63104** | YWHAZ | 14-3-3 protein zeta/delta (Protein kinase C inhibitor protein 1) | 245 | 0.9018 | 517 | 8282 | No |
| **P78362** | SRPK2 | SRSF protein kinase 2 (EC 2.7.11.1) (SFRS protein kinase 2) | 688 | 0.9278 | 441 | 8161 | No |
| **Q03001** | DST | Dystonin (230 kDa bullous pemphigoid antigen) | 7570 | 0.9152 | 58 | 7806 | Yes |
| **Q5VST9** | OBSCN | Obscurin (EC 2.7.11.1) | 7968 | 0.9118 | 14 | 7769 | Yes |
| **P38398** | BRCA1 | Breast cancer type 1 susceptibility protein (EC 2.3.2.27) | 1863 | 0.9160 | 456 | 7655 | Yes |
| **Q15149** | PLEC | Plectin (PCN) (PLTN) | 4684 | 0.8815 | 138 | 7611 | Yes |

Entry is the UniProt entry of the protein; Gene is the preferred gene names used in the literature; Length denotes to the length of the protein sequence; # of ints is the number of interactions adapted from PICKLE database; Disease is represented as a dichotomous variable adapted from UniProt's 'Involvement in disease' column.

## Discussion

### Proteome evolution during ages

As shown in the Methods section, relative frequencies of residues in a protein decide on its CAIR. Obviously, biochemical properties of amino acids play a central role in determining their primary relative frequencies in *de novo* proteins. Such dissimilarity of chemical properties would encourage unbalanced primary relative frequencies, thus lesser CAIRs, as shown in our results for younger proteins in Figure 4D. This finding follows the theory of *de novo* gene birth from non-coding DNA (Neme et al., 2017; Wilson et al., 2017). Nevertheless, during the course of evolution, residues are subjected to random mutation which equalizes their relative frequencies. This, in turn, increases the CAIR of proteins as they age which also agrees with the trend of CAIRs in Figure 4D. This can be a corroborating rationale for the study carried out with a different methodology in which it is shown that intrinsic disorder of proteins negatively correlates with gene age (Banerjee & Chakraborty, 2017). Random mutations aside, natural selection’s bias in favour of more complex proteins may have also contributed to increasing the CAIR in older proteins. This is also noticeable from the results seen in Figure 2 verifying the identical behaviour of natural selection toward all living organizations. Accordingly, it is not surprising that the human proteome includes a negatively-skewed distribution whose lesser CAIRs are mostly associated with proteins expressed by younger genes. Unfortunately, the literature lacks any thorough investigation on linguistic complexity of proteins and gene ages.

Moreover, it is evident from Figure 4C that the younger eukaryotic proteins are shorter than their older counterparts. A significant decline is noticed, however, in the length, interactions, and mutual information of proteins during the old ages. This refers to the different evolutionary rates of prokaryotic and eukaryotic genes and is compatible with the findings of previous studies (Alba & Castresana, 2005; Wolf et al., 2009). Even for proteins of prokaryotic ages, the trend of increase in protein length and interactions is observed; nonetheless, the trend line is discrete from that of eukaryotic proteins. Interestingly, this decline is not seen in Figure 4D, as the CAIR, in both eukaryotic and prokaryotic settings, is identically affected by directional selection. The dome shape increase of disease indicators from younger to older proteins refutes the justification of the study by Elhaik et al claiming that the slower rate of evolution in older genes is ‘an artefact of increased genetic distance (Elhaik et al., 2006)’.

Furthermore, it is demonstrated in Figure 4B that interactions increase as genes age which agrees with the literature (Saeed & Deane, 2006). However, the rate of increase seems to be slower in a prokaryotic setting. That means the rate of interaction turnover in eukaryotic proteomes may be comparatively higher. This might be due to the denser networks of eukaryotes and the gene duplication (Wagner, 2003). It is also noteworthy that the rate of interaction turnover seems to be non-decreasing as eukaryotic genes get older. Lastly, mutual information might be considered as an assembly of protein information and interactions. So as seen in Figure 4A, the trend of EMIP is also increasing by time for both eukaryotic and prokaryotic genes.

### Dynamics of protein networks

A protein network is considered ‘stable’ when the odds of network malfunction is tiny. Among various protein networks, particular ones malfunction often, and they generally are responsible for the networks of non-communicable diseases. According to the theorem presented in the Methods section, the number of interactions is negatively correlated to network stability. This deduction is contrary to what is seen in Figures 4B, 5B, and the literature (Jonsson & Bates, 2006; Oti et al., 2006; Xu & Li, 2006). According to the literature, the PPI networks of disease genes are different in topology containing more interactions, as it has been similarly shown in the mentioned figure panels. Previously, it was clarified that the interactions increase with a stationary rate as a gene ages. As a matter of fact, there are also many genes encoding for proteins of the cellular organisms that have not established any interactions. Hence, a confounding factor that might be overlooked would be the primary stability of the protein itself. In other words, proteins that are prone to malfunction need substantial interactions to compensate for their malfunctions within their network. That is the reason for increasing interactions along with the disease occurrences. In the literature, the finding of the increased mutation rate of disease genes may reason their increasing interactions (Smith & Eyre-Walker, 2003). Natural selection might be the other reason which would favour interactions merely for faulty networks rejecting in others.

As mentioned previously, an inaccurate estimate of a protein’s stability would be its CAIR. Figure 4D illustrates that CAIRs of disease proteins are significantly higher than those of non-disease proteins. However, the incongruous decrease of CAIRs in rare disease comparing with extremely rare diseases had not expected in the inferences of our theorem. This might be, in part, the influence of abundant younger proteins responsible for rare diseases that have not developed interactions yet. Proteins with less CAIR are more stable, yet they are negatively selected as they do not contain sufficient information. Complexity allows a protein to have the potential capacity to carry more intricate functions (Babu, 2016). Thus, in order to obtain higher functionalities, proteins grow both in their sizes and CAIRs. Consequently, this necessitates new interactions to arrive. However, new protein interactors may take millions of years to appear and reform the instability of the network (Fraser et al., 2002). These cycles begin all over again as the new proteins come into existence. All these aside, the essentiality of a protein’s function is an argument that should not be dismissed. Generally, old proteins are more crucial to forming life than younger ones (Chen et al., 2012). The less crucial roles of younger proteins render their diseases to be less severe. As human ages, numerous interactors in various networks are cancelled out causing these networks to malfunction. The coincidental rush of diseases at the late ages of human life may be caused by the accumulative effect of interactors removal and the loss of proteostasis (Kaushik & Cuervo, 2015; Labbadia & Morimoto, 2015). So, even substantial interactions cannot fully guarantee networks of complex unstable proteins.

### Gene age, hubs, and human diseases

Our results presented in Figure 5C illustrates the critical role of proteins encoded by the older genes to be responsible for a broad spectrum of diseases. This finding is in total agreement with a previous study in the literature highlighting the importance of ancient genes in human genetic disorders (Domazet-Lošo & Tautz, 2008). We also discussed that the older genes might take the leading role in creating the necessary fundamentals of life. A series of papers by Barabási et al dedicatedly demonstrate the distinction between disease genes and the essential genes. According to their work, disease genes are in most cases non-essentials being located at the periphery of the network, rather than being a central hub (Barabási et al., 2011; Domazet-Lošo & Tautz, 2008; Goh et al., 2007). In Figure 5D, although the trend of the increase in CAIR is parallel to more odds of disease to occur, the proportion of rare to extremely rare diseases suggests that more prevalent diseases are not associated with the proteins with the highest CAIRs. This is also evident from the overall comparison of CAIRs in disease occurrence categories in Figure 4D. This finding puts forward an argument that the most complex proteins located at the centre of networks are encoded by the essential old genes that in case of their malfunction, the condition would be fatal or cause an extremely rare and severe disease. However, the more prevalent diseases are caused by comparably less complex proteins that are indeed younger than central hubs. So, the proportion of rare to extremely rare diseases in Figure 5 is of great importance and should be noticed as they agree with the scenarios presented by Barabási et al.

### EMIP and human diseases

A disease may occur when the process of network compensation works inaccurately. For any network, the removal of the nodes would weaken its stability. This fact is evidently derived from our theorem. As we discussed, it is the reason why many of the unstable networks have urged to have many interactions. Integrating the effect of interactions with the protein information is what mutual information tries to demonstrate. In a sense, the mutual information of a protein is the information that is identical between the protein and its network. Thus, in the case of a malfunction, it would not result in a disease, as the network already carries the identical information. Estimation of mutual information, mathematically, is equivalent to the difference between the information of the network as a whole and the scalar summation of information that interactors carry when they are not interacting within a network. This, of course, will show how much capacity the network carries to compensate for its malfunction. In Figures 4A and 5A, the superior relation between EMIP and disease occurrence categories stems from this fact.

### On the existence of diseases

Based on what is discussed above, the scenario of a gene to cause a disease or not is being summarized as the following (Figure 7). When the evolution was in its initial periods, the involved proteins for life network was mainly the crucial ones. The metabolic pillars of life owe the fundamental and critical components to the first formation of these networks. There are two types of proteins at this stage, i.e. either ‘robust’ or ‘weak’. To define, robust proteins are those which are structurally not very susceptible to malfunction. These proteins have evolved without many interactions comparing to others. Hence, during the path of evolution, and still, we cannot detect as many interactions for them. On the other hand, weak proteins would have caused vital errors that result in severe diseases. It would be reasonable to assume that the incidence of such diseases would have been higher at ancient ages. So we may expect to observe many interactions of them till now. According to our theorem, it can be reasoned that these interactions would bypass the hub protein in case it malfunctions. So the incidence of such diseases would have been drastically lessened by now. As a general point, for an unstable weak protein that initiates a network, there are two prospects to be considered, i.e. being able to develop a mature network at the time of the investigation, or still getting involved in an immature one. By definition, mature networks would be those which have totally cancelled out the adverse outcomes of the malfunctioning hub protein. Based on the theorem, maturity is an ideal that no network can satisfy it in a biology setting.

**Figure 7.**
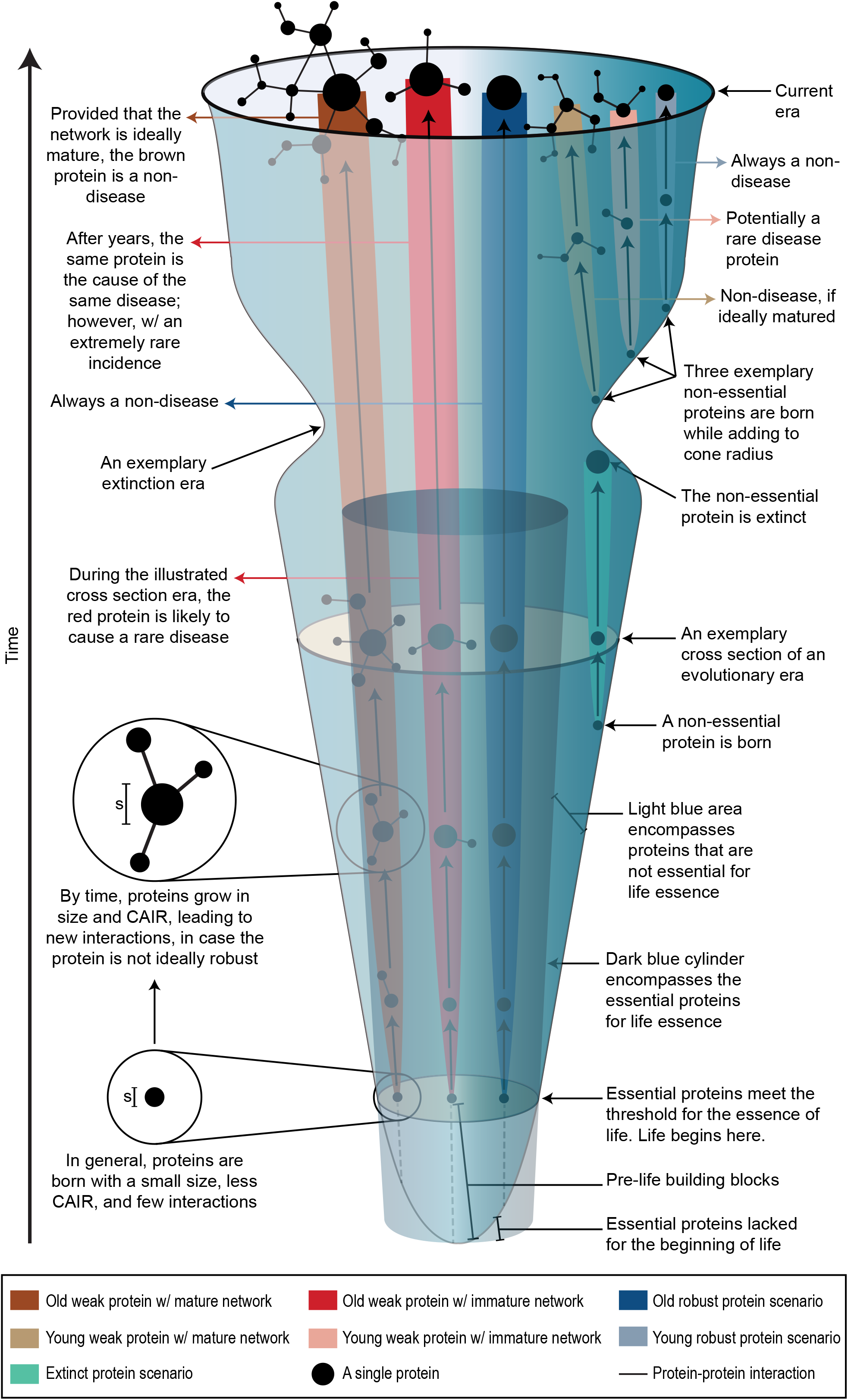
The cone of life and the evolution of diseases. The cone of life summarizes the probable scenarios for proteins put forward in the discussion section. It is to be noted that the shape of the cone is schematic and an exemplary instance of every possible occasion has been schematically illustrated. Also, for the clearness of the drawing, the dark blue cylinder has been cut in half in order not to block other elements. The diameter of the cone in any cross section shows the amount of existing protein material in that given time period. The proteins are born from the circumference of any cone base. As shown in the figure, a general rule would be that diseases are a result of weak proteins with immature networks. Details have been depicted in the figure for every protein scenario. It is to be highlighted that the eukaryotic and prokaryotic proteomes have not been discerned in the figure. s, size; w/, with.

Having mentioned all these about the archaic proteins, it should also be noticed that the proteins which have come into existence in more chronologically proximal periods would have by far lower chances of being a vital hub. The functioning network of life, in one piece, has less to do with a mammalian protein than an archaic cellular respiration protein. So, the other category of proteins is that of the contemporary period on which the logical assumption would be that they are either non-hubs or if hubs do not function in places crucial to life. The contemporary category, i.e. the young proteins, again, is subdivided into robust and weak proteins. Robust young proteins are those without interactions which are not very much susceptible to cause diseases, because if otherwise, evolution would bring interactions for them. It is noteworthy that very young proteins are mainly very stable, simple, and proteins with minor functional capacities. Stable young proteins may by time change to unstable complex proteins because of the random mutations and the natural selection’s bias toward more complex proteins. In the way of transformation, most of the weak young proteins arise which are primarily responsible for the prevalent metabolic diseases of the current evolutionary era. Since the average evolutionary rate interaction turnover has not satisfied the optimum number of network nodes for them, they are apprentices susceptible to malfunction. It would be easy to infer that these proteins cause less severe diseases comparing to archaic proteins, as their functions are still not vital. Nevertheless, they cause diseases with higher incidences.

### Applicability of the method

The methodology we have presented in the next section is not sensitive to the level of taxonomy, i.e. whether the calculation is for a species, a genus, an order, a kingdom, or the complete tree of life. The reason for this fact is that calculating the amino acid frequencies is the same considering a species proteome or any other more extensive proteomic combination of taxonomic levels. Also, we can calculate the Shannon entropy for a single protein, or a peptide. This insensitivity to the size of the network in calculation enables a homogenous analysis through the whole tree of life.

### A perspective of future studies

Future studies may focus on each of the non-communicable diseases to elaborate more on the speculations made in this study. Communicable diseases may also be the focus of further studies to investigate the CAIRs of organisms and probable relations to their pathogenicity. Treatments that are targeting networks with high-CAIR interacting protein crowds would be an option to be explored. Moreover, better estimations for mutual information would be of great interest. Besides, the notation of the wave of life and CAIR comparisons may bring further arguments to taxonomists. Lastly, the theorem may be used in various fields of science which are shaped by networks.

## Methods

### Introducing the Calculated Average Information per Residue (CAIR) and the Protein Information (PI)

Proteins, form a mathematical point of view, are once randomly-occurred sequences of residues that have gone through a process of selection in nature. Considering this fact, a protein structure can be defined using a random sequence that carries mathematical information. The information that a protein carries may be defined as to be equivalent to the amount of uncertainty in predicting its residues. It is to be highlighted that regardless of the protein conformation, the information of a protein is determined merely by its primary structure. The average information carried by a residue in a protein is calculated by Shannon’s entropy (*H*) equation as below:

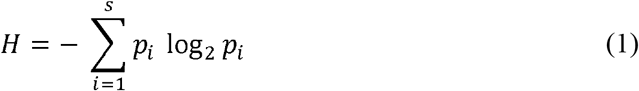

where *p_i_* is the probability of state *i*, and *s* is the total number of possible states. In the current context, the CAIR notion is introduced to be the same as Shannon’s entropy except for the logarithm base which is 22 in the former and 2 in the latter. In other words, CAIR is the 22-ary of Shannon’s entropy and is formulated as:

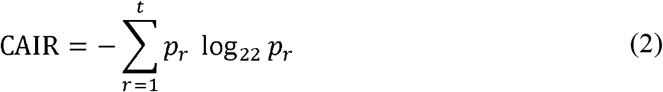

in which *r* is a numeral given to each residue, *t* is the total number of residues, *p_r_* is the relative frequency of *r*^th^ residue in the protein. More simply, the CAIR could be written as:

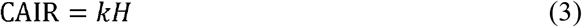

in which *H* is Shannon’s entropy in equation (1), and *k* is a constant equivalent to:

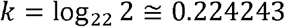

As it is evident from equation (3), the patterns in the results obtained in the current article were independent of the base of the logarithm; however, the scale of entropy would be much more tangible considering the base of 22, as the ideal proteinogenic alphabet contains 22 letters. Deriving from CAIR, protein information (PI) is the amount of information carried by all the residues of a protein as of the following equation:

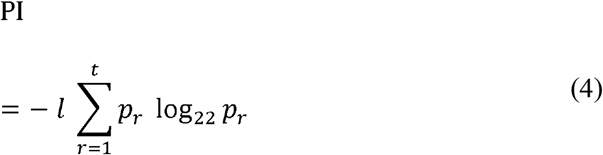

in which the variables are the same as those in equation (2), and *l* is the length of the protein. The notations of PI and CAIR are proposed, instead of the conventional *H*, for the fact that they are more expressive and pertinent for the field of proteomics. It would also be humbly proposed – with an analogy to Shannon’s bit – to use ‘pit’, i.e. protein unit, for the unit of CAIR, PI, and EMIP in order to be readily comprehensible. As an example, one kilo-pit would be equivalent to the PI of a 1000-lengthed protein whose residues have equal frequencies.

### Mathematical Theorem

Suppose {*X*},{*Y*_1_},{*Y*_2_}…, {*Y_n_*} are sets of sequences from which {*Y*_1_},{*Y*_2_}…,{*Y_n_*} are all dependent to {*X*}, but are pairwise independent from each other, not necessarily identically distributed random variables having characteristic functions of *φ*_1_, *φ*_2_,…, *φ_n_*, distribution functions of *f*_1_,*f*_2_,…, *f_n_*, and entropies of *H*_1_, *H*_2_,…, *H_n_*. Let Φ be the characteristic function of {*X*}, 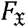 be its distribution function, and 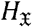 be its entropy. Then:

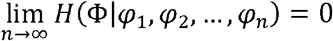

***Proof.*** According to the definition of mutual information, we can write the following relations (Cover & Thomas, 2012):

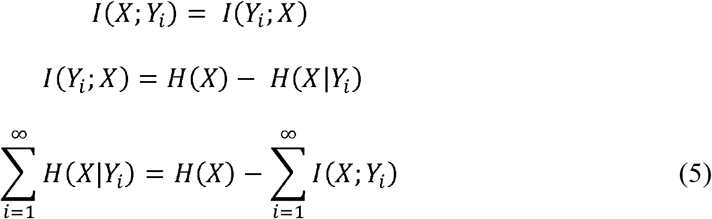

Corollary. Non-negativity of mutual information(Cover & Thomas, 2012):

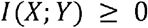

with equality i☐ *X* and *Y* are independent.

Based on the corollary of non-negativity of mutual information, and because *Y_i_* are all dependent on *X*, the mutual information of *X* with respect to all *Y_i_* is always positive:

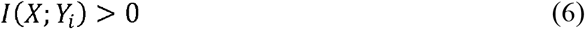

from which it can be inferred that infinite sum of positive values yields not to infinite, but to the maximum mutual information possible, i.e., the entropy of *X*:

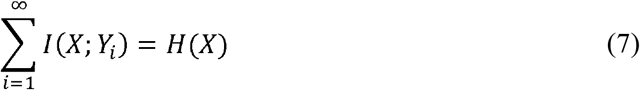

So, substituting equation (7) in equation (5):

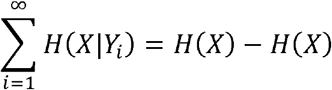

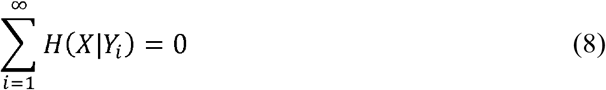

Also, since 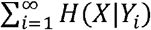 is the same as 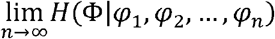, thus:

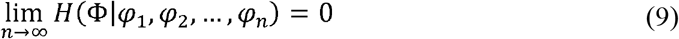

### Biological inferences and hypotheses

In the above theorem, supposing Φ be a protein with the amino acid sequence of {*X*}, interacting with *n* number of other proteins, namely *φ*_1_,*φ*_2_,…, *φ_n_*, with sequences of {*Y*_1_},{*Y*_2_}…, {*Y_n_*}:

1. Said interactions are mathematically interpreted as the dependency of 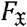 on *f*_1_,*f*_2_,…,*f_n_*. To elucidate, the probability distribution functions of proteins correspond to their Boltzmann distributions. Because of the induced-fit nature of biochemical interactions, it might be plausible to consider the distribution functions to be dependent on one another. For that reason, each interaction is deduced as the dependency of two distributions in the theorem.
2. As the theorem suggests, 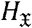 indicates Shannon’s entropy of the protein Φ with an amino acid sequence of {*X*}. This might be interpreted as the extent of probable variations that could lay in the primary structure of a protein affecting its function. This measure would be an inaccurate estimate of a protein’s malfunction as no biochemical conditions have been taken into account. Despite its inaccuracy, we have included the CAIR as an indicator of diseases in our analysis. The intuitive hypothesis would maintain that the CAIR is significantly more in disease proteins than non-disease ones for their surplus odds of having potential disadvantageous variations in the primary structure leading to malfunction and disease.
3. According to the assumptions of the theorem, the notation of ‘information’ could also apply to any system containing proteins, i.e. a particular metabolic pathway, an interactome, diseasome, or the entire living organisms. Measuring the information is independent of the size of the network. This property allows us to calculate the information among the different taxonomic hierarchies and to utilize the results in practice. Also, in a single organism like *Homo sapiens*, it would allow us to compare different disease pathways, track hub proteins, and discern potential disease proteins from non-disease ones.
4. The length of a protein sequence determines the PI, as shown in equation (4). We have not included the PI itself as an indicator in the study, but have included both the CAIR and the protein length separately. Proteins with longer sequences carry more information and are more prone to malfunction as the overall odds of a faulty residue in their sequence is higher compared to a protein with a shorter sequence.
5. Considering equation (9), it is understandable that the conditional entropy of Φ with respect to the knowledge of *n* number of interactors, i.e. *φ*_1_,*φ*_2_,…,*φ_n_*, would equal zero when *n* approaches infinity. The conditional entropy designates the new information carried by the Φ protein when functioning in its network with other proteins of *φ*_1_ to *φ_n_*. So, as a preliminary and naive inference, it could be easily inferred from the theorem that networks with more interactions are more stable. Therefore, we have included the number of interactions as an indicator of human diseases to test our hypothesis.
6. Although equation (9) is a relatively straightforward approach to calculate the stability of a network, it can be shown that its exact quantitative calculation is not possible in proteomic analysis. An equivalent measure of network stability would be to use equation (6) in which mutual information is shown to be always positive. Unlike the conditional entropy which is negatively correlated to the network stability, the mutual information is positively correlated. To quantitate mutual information in the current context, we propose the following estimation, henceforth referred to as EMIP (*μ*):

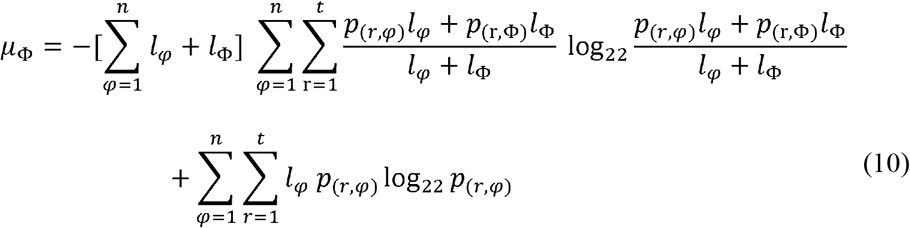

in which *μ*_Φ_ is the mutual information for the Φ protein, *n* is the number of interactions, *l_φ_* is the length of the *φ*^th^ interactor, *l*_Φ_ is the length of protein Φ, *r* is a numeral given to each residue in proteins, *t* is the total number of residues, *p*_(*r*, *φ*)_ is the relative frequency of *r*^th^ residue in *φ*^th^ interactor, and *p*_(*r*, Φ)_ is the relative frequency of *r*^th^ residue in Φ protein. EMIP has also been included as an indicator of diseases in our analysis.
7. An essential element in the course of life evolution and human disease analysis is to consider the gene age of proteins. It is conceivable to hypothesize that the chronological data of genes can be very much associated with the indicators of human diseases. One reason is the additive effect of gene ages to introduce new interactions. The second reason is based on the assumption that natural selection can show bias towards proteins with a specific range of CAIRs. Also, the third reason is the possibility that the older proteins can grow to have longer sequences during evolution. Therefore, we have also included the gene ages as a covariate of human diseases in our analysis.
8. According to the theorem, it is also inferred that in an ideal condition with an infinite number of interactors for a protein, the functionality of such a hub protein reaches an infallible state, i.e. no disease would ever happen. This could also be applied to other fields of science to reason why there is an order in complex systems with innumerable components. The rate of decline in error in our theorem as shown in equation (9) under random conditions is generally consistent with the simple statistical rule of 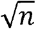 as proposed in physicist Schrödinger’s ‘what is life’ (Schrödinger, 1944).

### Protein database

For taxonomic comparisons, all complete proteomes were extracted from the UniProt (Consortium, 2019) FTP server, freely available at *ftp://ftp.uniprot.org/pub/databases/uniprot/current_release/knowledgebase/complete/*. Both SwissProt and TrEMBL files were downloaded in FASTA format. We calculated protein entropies separately for each entry in both files including a total of more 61 billion amino acids and merged them with every individual organism. Then, we used the calculations for further statistical analyses.

Besides, for human proteins analyses, the complete list of *Homo sapiens* proteins was downloaded directly from the UniProt website in a tab-separated (.tab) format containing the following columns: ‘Entry’, ‘Length’, ‘Sequence’, ‘Orphanet’, and ‘Involvement in disease’. Data were updated on 22^nd^ April 2020 with UniProt release 2020_02.

### Protein-protein interactions database

Protein-protein interactions in human proteome were obtained from Protein InteraCtion KnowLedgebasE (Gioutlakis et al., 2017) (PICKLE) meta-database, the release of 2.5. PICKLE is a cross-checked integration of all available human PPIs included in BioGRID, IntAct, HPRD, MINT, DIP databases. The default filter mode was selected to download 191,113 binary interactions among 16,418 UniProtKB/SwissProt entries. All interactions were included in our study for further analysis.

### Taxonomy database

Taxonomy data of organisms were also extracted from the UniProt FTP, the release of 2020_02. All organisms were included and matched according to their ‘OX’, i.e. organism number used by UniProt and other databases. The evolutionary tree was plotted based on a landmark study (Hug et al., 2016) by Hug et al, published in 2016, with a review of updates (Carnevali et al., 2019; Carr et al., 2019; Cavalier-Smith et al., 2014, 2018; Dombrowski et al., 2019; Eloe-Fadrosh et al., 2016; Eme et al., 2017; Hahnke et al., 2016; Hamilton et al., 2016; Jay et al., 2018; Jungbluth et al., 2017; Kevbrin et al., 2020; Kirkegaard et al., 2016; Martinez et al., 2019; L. Momper et al., 2017; L. M. Momper et al., 2018; Munoz et al., 2016; Pavan et al., 2018; Y. Wang et al., 2019; Ward et al., 2019; Youssef et al., 2019; Zhou et al., 2020) since then until April 2020. All updates were added to the tree and were matched with UniProt taxonomy data.

### Diseases database

According to cross-references between UniProt and Orphanet, related epidemiological data were downloaded and extracted from the Orphanet database (Weinreich et al., 2008) in XML format. The file was then used in Python code for further analysis. Only diseases with at least one reported worldwide occurrence were included. Normally, in the Orphanet database, occurrences are reported from one or more of the following categories: annual incidence, cases/families, lifetime prevalence, point prevalence, and prevalence at birth. In the case of more than one reported occurrence, the priority of selection was for incidence, prevalence at birth, point prevalence, respectively. Accumulative occurrence data, i.e. cases/families and lifetime prevalence, were excluded from the analysis.

### Analysis of taxonomic hierarchies

The following steps were carried out to implement the theorem over the taxonomic hierarchies (Figure 8A):

1. TrEMBL and SwissProt FASTA files of complete proteomes were downloaded from UniProt KnowledgeBase. A total number of ~180 million protein entries were included.
2. The frequencies of all amino acid residues were calculated with regards to all protein entries.
3. All proteins were grouped according to their organisms using organism IDs.
4. Organisms that are not proteomes were excluded. Duplicates are also removed.
5. Viruses are excluded from the study.
6. Shannon’s entropy was calculated according to the residue frequencies for all included non-virus proteomes.
7. Taxonomic data were downloaded directly from the UniProt website.
8. The most updated tree of life was drawn after a thorough review of the literature until April 2020.
9. Organisms were grouped with respect to taxonomic hierarchies.
10. Organisms with unknown taxonomic lineage were excluded from the analysis.
11. Brunner-Munzel test was performed for every bifurcation node through the tree of life. The level of significance was 0.05. Preparation of data and protein information calculations were all executed in Python 3.8.0 (Van Rossum & Drake, 2009) using NumPy (Oliphant, 2006), Pandas (McKinney & others, 2010), and Biopython (Cock et al., 2009) libraries. Statistical analysis and violin plots were carried out using R-3.6.0 (R Core Team, 2019) with brunnermunzel (Ara, 2020) and Plotly (Sievert, 2018) packages.

**Figure 8.**
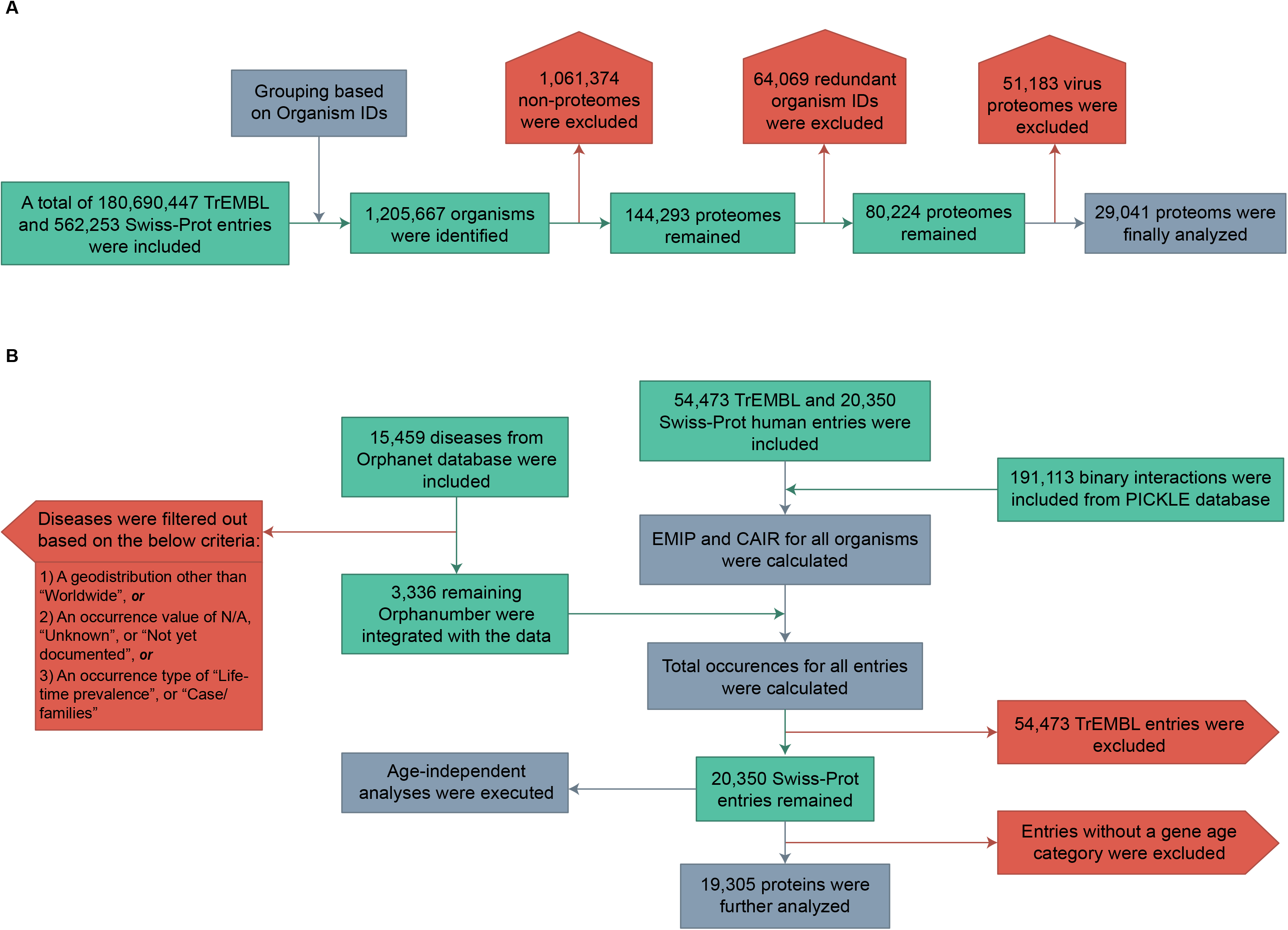
Detailed steps needed to carry out the presented methodology. **(A)** Flowchart shows how the proteins were included, grouped according to their respective organisms, and the exclusion criteria. **(B)** A similar flowchart explaining the steps used to integrate human proteome data, PICKLE interactions, and Orphanet diseases. Green rectangles, steps; purple rectangles, executions; red flags, exclusions.

### Analysis of human disease proteins

The following steps were carried out to implement EMIP and analyse disease occurrence categories (Figure 8B):

1. The human proteome was downloaded from the UniProt database with the organism ID of 9606. 20,350 of reviewed and 54,473 of unreviewed proteins were included in the study.
2. CAIR was calculated for all human entries using the sequences of residues.
3. Protein-protein interactions were extracted from the PICKLE database as the default UniProt normalized file with a total number of ~190k interactions.
4. Interactions were altered in order to match the UniProt ‘interactions with’ column. This was done to keep the homogeneity of the data.
5. EMIP was then calculated for all entries with the help of the PPIs.
6. The unreviewed proteins were excluded after the calculations of disease indicators because they have not been reported to cause any diseases.
7. Ordinal age categories of genes were merged to the file using the consensus data article (Liebeskind et al., 2016).
8. Disease categories are ranked using R into three groups of no diseases, extremely rare diseases, and rare diseases.
9. Disease indicators were also categorized into eight groups to draw Likert plots.
10. Statistical differences between groups were tested among variables with the DTK test. Significance levels of the DTK test were added to the comparison graphs with stars. The highest significance level was set to 0.05. Graphs of comparisons were plotted in R with DTK (Lau, 2013) and ggpubr (Kassambara, 2019) packages. Likert plots were plotted with HH (Heiberger, 2019) package. Networks were plotted using igraph (Csardi & Nepusz, 2006) package.

## Data availability

Data used for analysis is available as the following files. FASTA files of all Swiss-Prot and TrEMBL entries are publicly available from UniProt’s FTP server at https://ftp.expasy.org/databases/uniprot/current_release/knowledgebase/complete/. Also, all non-redundant proteomes could be downloaded from UniProt website: https://www.uniprot.org/proteomes/?query=redundant:no&format=tab&force=true&columns=id,name,organism-id,lineage&compress=yes. Tab-separated format of human proteome data used in our analysis is achievable from https://www.uniprot.org/uniprot/?query=proteome:UP000005640&format=tab&force=true&columns=id,reviewed,genes(PREFERRED),protein%20names,sequence,database(Orphanet),comment(INVOLVEMENT%20IN%20DISEASE),interactor&compress=yes. Additionally, PICKLE interactions are freely available from http://www.pickle.gr/Data/2.5/PICKLE2_5_UniProtNormalizedTabular-default.zip. Orphanet data is also freely available from http://www.orphadata.org/data/xml/en_product9_prev.xml. Data of gene ages are adopted from the Gene-Ages GitHub repository at https://github.com/marcottelab/Gene-Ages/raw/master/Main/main_HUMAN.csv. Supplementary information is available in the online version.

All Python and R codes necessary to reproduce all parts of the analysis and for the illustration of the figures are available under the MIT license on our GitHub repository at https://github.com/synaptic-proteolab/CAIR_EMIP, or the Zenodo link at https://zenodo.org/record/3975465. For executing codes online on cloud servers, Google Colab links are also available on the GitHub page. Python (https://www.python.org/downloads/), Jupyter Notebook (https://jupyter.org/install), R (https://cran.r-project.org/), and RStudio (https://rstudio.com/products/rstudio/download/) are all freely available for the public.

## Acknowledgments

Authors would like to thank Dr Saeed Sadigh-Eteghad, PhD in neurosciences, Tabriz University of Medical Sciences, and Dr Pedram Dindari, a PhD candidate in computer sciences, University of Tabriz, for their assistance during the preparation of the manuscript. No funding to declare.

## Author contributions

FK proposed the mathematical theorem and its proof. SD gathered and handled the large data and prepared the data for analysis using Python. FK reviewed the literature to illustrate the tree of life. FK analysed the data using R. SD and FK prepared the codes for open-source publishing. FK wrote the manuscript, and SD agreed with all sections. FK designed the figures and SD prepared the tables of the manuscript.

